# Coordinated Changes In The Accumulation Of Metal Ions In Maize (*Zea mays ssp. mays L.*) In Response To Inoculation With The Arbuscular Mycorrhizal Fungus *Funneliformis mosseae*

**DOI:** 10.1101/135459

**Authors:** M. Rosario Ramirez-Flores, Ruben Rellan-Alvarez, Barbara Wozniak, Mesfin-Nigussie Gebreselassie, Iver Jakobsen, Victor Olalde-Portugal, Ivan Baxter, Uta Paszkowski, Ruairidh J.H. Sawers

## Abstract

Arbuscular mycorrhizal symbiosis is an ancient interaction between plants and fungi of the phylum Glomeromycota. In exchange for photosynthetically fixed carbon, the fungus provides the plant host with greater access to soil nutrients via an extensive network of root-external hyphae. Here, to determine the impact of the symbiosis on the host ionome, the concentration of nineteen elements was determined in the roots and leaves of a panel of thirty maize varieties, grown under phosphorus limiting conditions, with, or without, inoculation with the fungus *Funneliformis mosseae.* Although the most recognized benefit of the symbiosis to the host plant is greater access to soil phosphorus, the concentration of a number of other elements responded significantly to inoculation across the panel as a whole. In addition, variety-specific effects indicated the importance of plant genotype to the response. Clusters of elements were identified that varied in a coordinated manner across genotypes, and that were maintained between non-inoculated and inoculated plants.

**Abbreviations:**

NC: non-colonized
M: mycorrhizal
SDW: shoot dry weight
ICP-MS: inductively coupled plasma mass spectrometry
PC: principal component

## Introduction

Plants require 14 essential mineral elements to complete their lifecycle, namely nitrogen (N), phosphorus (P), potassium (K), sulfur (S), calcium (Ca), magnesium (Mg), iron (Fe), manganese (Mn), copper (Cu), zinc (Zn), molybdenum (Mo), boron (B), chloride (Cl) and nickel (Ni) (Marschner, 2012). Depending on their concentration in the plant, these elements can be classified as macronutrients or micronutrients. In addition, elements can be classified into four major groups based on their requirement for 1) synthesis of biomolecules, 2) energy transfer, 3) ion balance and 4) electron transport. As a by-product of nutrient and water acquisition, plants will also take up a number of additional non-essential elements that may, in high concentrations, be toxic, such as Aluminum (Al), Arsenic (As), Cadmium (Cd), Cobalt (Co), Selenium (Se), Strontium (Sr) and Rubidium (Rb). A deficiency of mineral elements has detrimental consequences on plant fitness or, in the agronomic context, crop yield, and plants have developed a number of strategies to promote uptake in nutrient deficient soils and optimize the efficiency of internal use. Such responses include modification of the root system architecture, induction of high affinity nutrient transporters, remobilization of internal resources, growth arrest, down-regulation of photosynthesis and the induction of senescence (Lynch, 1995; Aibara and Miwa, 2014; Whitcomb et al., 2014). In addition, plants form mutualistic associations with rhizosphere organisms, such as nitrogen-fixing rhizobia or arbuscular mycorrhizal (AM) fungi (Bucher, 2007; Parniske, 2008; Vance, 2014).

AM symbiosis is a mutualistic interaction established between soil fungi belonging to the phylum Glomeromycota and the majority (70-90%) of land plant species (Schübler *et al.*, 2001; Smith and Read, 2008). The capacity to form AM symbiosis has been retained in the major cereal crop species throughout domestication and improvement *(e.g.* Koide *et al.*, 1988; Hetrick *et al.*, 1992; Kaeppler *et al.*, 2000; Sawers *et al.*, 2008), as has a conserved molecular machinery required for symbiotic establishment and nutrient exchange *(e.g.* Paszkowski *et al.*, 2002; Gutjahr *et al.*, 2008; Yang *et al.*, 2012; Willman *et al.*, 2013; Liu *et al.*, 2016; Nadal *et al.*, 2017). One of the major benefits of AM symbiosis to the plant host is enhanced nutrient uptake as the result of enhanced soil foraging by an extensive network of root-external fungal hyphae (Bago *et al.*, 2003; Finlay, 2008). An increase in P uptake in P limiting soils is well established as the primary physiological consequence of AM symbiosis on the plant host (Bucher, 2007). AM symbiosis, however, has a broad impact on host mineral nutrition, potentially increasing the uptake of additional essential nutrients such as N, Cu, Fe, Mn and Zn or limiting the uptake of potentially toxic elements such as Cd, Pb, Hg and As (Govindarajulu *et al.*, 2005; Jin *et al.*, 2005; Göhre and Paszkowski, 2006; Guether *et al.*, 2009). These effects may be mediated through direct transport by the fungi, or by alterations in root system architecture and physiology. Transcriptomic and functional analyses have identified plant encoded nutrient transporters specifically expressed or up-regulated in mycorrhizal plants, including transporters of phosphate *(e.g.* Harrison *et al.*, 2002; Paszkowski *et al.*, 2002; Liu *et al.*, 2016), ammonium (Koegel *et al.*, 2013), sulfate (Giovannetti *et al.*, 2014) and sodium (Porcel *et al.*, 2016). To better understand the impact of AM symbiosis on plant nutrition, it is informative to consider the ionome - the total element composition - as a whole, investigating the relationships that exist between elements as a result of their interaction in soil chemistry, common uptake machinery, and the mechanisms of plant internal homeostasis (Baxter *et al.*, 2008; Baxter, 2015).

Here, the concentration of nineteen elements was determined in the leaves and roots of a panel of thirty maize lines, consisting of the 26 parents of the nested association mapping (NAM) population (McMullen *et al.*, 2009) and a number of additional varieties, grown in the greenhouse under P limiting conditions, with or without inoculation with the AM fungus *Funneliformis mosseae.* Maize is not only a crop of great agronomic importance, but also a model system well supported with genetic and genomic tools. Previous studies have shown commercial maize varieties to be well colonized by AM fungi and to typically present a strong growth response to AM inoculation under P limiting greenhouse conditions (e.g. Kaeppler *et al.*, 2000; Sawers *et al.*, 2017), while reverse genetics approaches are defining the importance of specific genes to the functioning of AM symbiosis in maize (Willman *et al.*,2013; Nadal *et al.*, 2017). The study panel represented a non-biased sampling of the broader genetic diversity of maize breeding lines, allowing generalization ad to the effect on specific elements, investigation of correlated responses across lines, and evaluation genotype specific responses.

Although P was the limiting nutrient in the experiment, the host ionome responded broadly to AM symbiosis, reinforcing the idea that chemical elements behave as a coordinated system when growth conditions are altered. Furthermore, the response among certain groups of elements was correlated, even when the response itself differed among maize varieties. Given interest in the agronomic application of AM symbiosis to increase the efficiency of fertilizer use, improve nutritional value, and maintain the concentrations of toxic metals at safe levels (Sawers et al., 2008; Fester and Sawers, 2011), these results provide a valuable reference dataset for further characterization of the effect of mycorrhizal colonization on the maize ionome under field conditions.

## RESULTS

### The maize ionome responds to inoculation with *Funneliformis mosseae*

To assess the impact of mycorrhizal colonization on the host ionome, root and shoot samples collected from a previously reported maize evaluation (Sawers *et al.*, 2017) were analyzed by inductively coupled plasma mass spectrometry (ICP-MS). In this experiment, 30 maize inbred lines, selected to maximize genetic diversity (McMullen *et al.*, 2009), were grown with (M) or without (NC) inoculation with *Funneliformis mosseae,* under phosphorus (P) limiting conditions (Sawers *et al.*, 2017). It was reported previously that plants were well colonized (about 60% of the total root length contained fungal structures, and about 30% of the total root length contained arbuscules) with an associated increase in shoot dry weight (SDW) of approximately two-fold (Sawers *et al.*, 2017). Here, in addition to the concentration of P included in the initial report, concentrations of a further 18 elements are presented. Initially, all genotypes were considered together, to generalize as to the main effect of fungal inoculation on the maize ionome (Fig.1; Table 1). Inoculation with *F mosseae* was associated with a significant increase in P concentration in both roots (Wilcoxon test, p < 0.05) and leaves (p < 0.01). In addition, in roots, a significant increase of Na (p < 0.001) and S (p < 0.05) was observed, along with a decrease of Cd, Co, Mn and Ni (all, p < 0.001). In leaves, there were significant increases in the concentration of Al (p < 0.001), Fe (p < 0.001) and a decrease of Zn (p < 0.05) concentration.

**Fig. 1.**
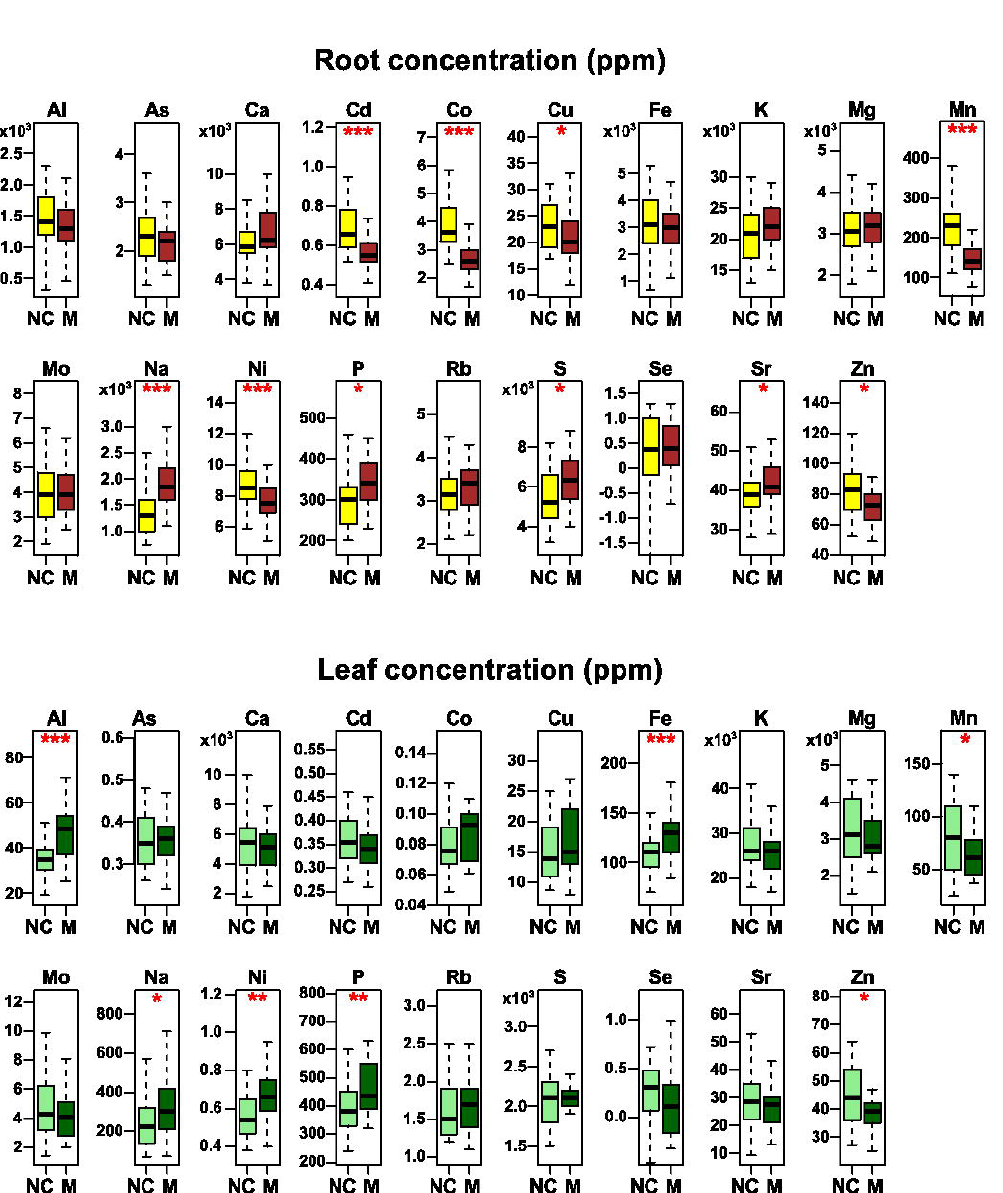
Element concentration responds to inoculation with *Funneliformis mosseae.* Concentration (ppm) of ninteen elements in the roots of non-colonized plants (yellow), the roots of colonized plants (brown), the leaves of non-colonized plants (pale green) and the leaves of colonized plants (dark green) determined by inductively coupled plasma mass spectrometry. Boxes show 1st quartile, median and 3rd quartile. Whiskers extend to the most extreme points within 1.5x box length; outlying values beyond this range are not shown. Ions for which accumulation differed significantly (Wilcoxon test) between NC and M plants indicated by * (p < 0.05), ** (p < 0.01) or *** (p < 0.001).

**Table 1.** Concentration of twenty ions in the roots and leaves of maize plants with and without inoculation with *Funneliformis mosseae.* Marginal mean (Conc; ppm) and standard error (SE) for the concentration of twenty ions in the roots and leaves of non-inoculated (NC) and inoculated (M) plants, calculated across thirty genotypes. r, Pearson correlation coefficient between NC and M plants. p, P-value from Mann-Whitney test for equivalent concentration in NC and M plants.

| Ion | Root |  |  |  |  |  | Leaf |  |  |  |  |  |
| --- | --- | --- | --- | --- | --- | --- | --- | --- | --- | --- | --- | --- |
|  | NC |  | M |  | r | p | NC |  | M |  | r | p |
|  | Conc (ppm) | SE | Conc (ppm) | SE |  |  | Conc (ppm) | SE | Conc (ppm) | SE |  |  |
| Al27 | 1500 | 100 | 1300 | 72 | 0.09 | 0.317 | 36 | 1.7 | 48 | 2.3 | -0.27 | 0.001 |
| As75 | 2.4 | 0.13 | 2.2 | 0.085 | 0.14 | 0.263 | 0.35 | 0.011 | 0.36 | 0.0094 | 0.46 | 0.539 |
| Ca43 | 6300 | 320 | 7000 | 380 | 0.12 | 0.112 | 5500 | 350 | 5100 | 290 | 0.6 | 0.615 |
| Cd111 | 0.68 | 0.02 | 0.56 | 0.014 | 0.36 | 0.000 | 0.36 | 0.01 | 0.35 | 0.0086 | 0.28 | 0.226 |
| Co59 | 3.9 | 0.15 | 2.7 | 0.099 | 0.7 | 0.000 | 0.085 | 0.0059 | 0.091 | 0.0059 | 0.5 | 0.160 |
| Cu65 | 24 | 0.97 | 21 | 1 | 0.53 | 0.036 | 15 | 0.83 | 18 | 1.6 | 0.32 | 0.173 |
| Fe57 | 3300 | 240 | 3000 | 150 | 0.14 | 0.684 | 110 | 4 | 140 | 6 | 0.08 | 0.001 |
| K39 | 21000 | 840 | 22000 | 670 | 0.56 | 0.260 | 28000 | 1400 | 26000 | 1100 | 0.49 | 0.343 |
| Mg25 | 3200 | 130 | 3100 | 110 | 0.71 | 0.976 | 3200 | 160 | 3000 | 120 | 0.57 | 0.491 |
| Mn55 | 240 | 17 | 140 | 6.8 | 0.42 | 0.000 | 82 | 6.3 | 64 | 3.6 | 0.7 | 0.047 |
| Mo98 | 4.1 | 0.31 | 4.1 | 0.21 | 0.6 | 0.796 | 4.9 | 0.49 | 4.5 | 0.44 | 0.8 | 0.631 |
| Na23 | 1400 | 91 | 1900 | 82 | 0.6 | 0.000 | 260 | 27 | 330 | 29 | 0.35 | 0.043 |
| Ni60 | 8.8 | 0.32 | 7.5 | 0.22 | 0.37 | 0.005 | 0.58 | 0.03 | 0.71 | 0.049 | 0.02 | 0.006 |
| P31 | 300 | 13 | 340 | 13 | 0.05 | 0.041 | 390 | 16 | 450 | 16 | 0.09 | 0.005 |
| Rb85 | 3.2 | 0.12 | 3.3 | 0.092 | 0.13 | 0.296 | 1.7 | 0.074 | 1.8 | 0.089 | 0.58 | 0.277 |
| S34 | 5500 | 250 | 6400 | 270 | 0.55 | 0.011 | 2100 | 53 | 2200 | 42 | 0.29 | 0.361 |
| Se82 | 0.59 | 0.24 | 0.48 | 0.15 | 0.19 | 0.906 | 0.31 | 0.089 | 0.018 | 0.11 | 0.22 | 0.086 |
| Sr88 | 40 | 1.2 | 43 | 1.6 | 0.22 | 0.042 | 30 | 2.2 | 26 | 1.6 | 0.74 | 0.407 |
| Zn66 | 85 | 3.8 | 74 | 3.1 | 0.21 | 0.032 | 46 | 1.9 | 39 | 1.2 | 0.63 | 0.017 |

### Patterns in ion concentration shift following inoculation with *F. mosseae*

To characterize relationships among ions, pairwise correlations were calculated among all 38 ion-inoculation combinations, separately for roots and leaves, across the thirty varieties evaluated (Fig. 2, 3). Correlation in the concentration of any given ion between NC and M plants ranged from 0.05 to 0.71 in roots and 0.03 to 0.74 in leaves (Table 1). Clustering of the correlation matrix revealed covariation among ions and between M and NC treatments (Fig. 2, 3). For certain ions M and NC treatments grouped together (e.g. K in both roots and leaves), indicating that patterns of variation among lines were maintained, while for other ions, M and NC treatments were not close in the clustering (e.g. P in both roots and leaves), indicating patterns of relative accumulation among lines to be changing under colonization. Clustering revealed also relationships among ions. The largest cluster was seen in the roots, consisting of Al, As, Co, Fe, Ni and Rb (Fig 2). Interestingly, this cluster is maintained under both NC and M treatments, although the correlations of the ions between NC and M treatments are low: *i.e.* the impact of AM symbioses on the concentration of these ions was genotype specific, such that level in NC plants did not well predict the level in M plants; and yet, whatever response did occur in a given variety was coordinated across the clustered ions.

**Fig. 2.**
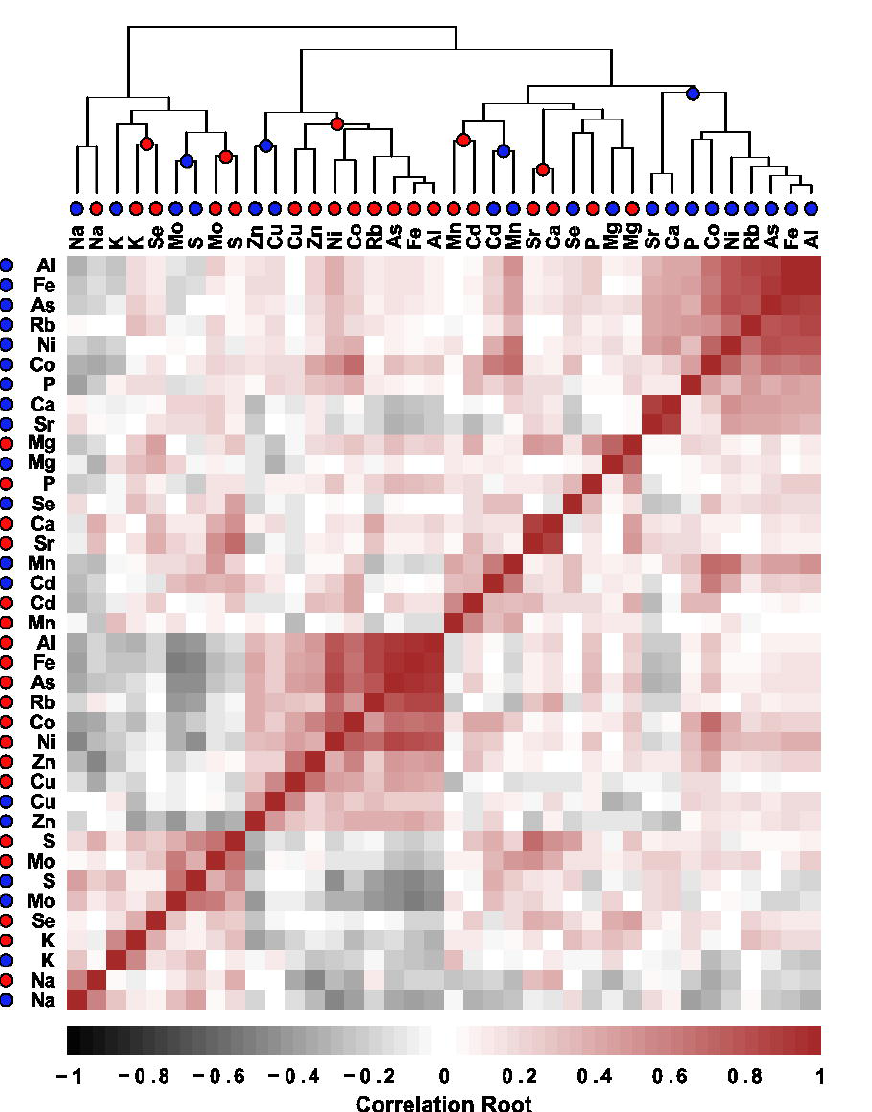
Patterns of ion concentration in roots shift following inoculation with *Funneliformis* mosseae. Pairwise correlation of ion concentration (colour-coded square) in the roots of thirty maize varieties grown with (indicated by red point adjacent to ion name) or without (indicated by blue point) inoculation with *F. mosseae*. The thirty-eight ion x inoculation combinations are clustered hierarchically. Nodes are marked with a red or blue point to indicate all adjoining lower order nodes to share the same inoculation status.

**Fig. 3.**
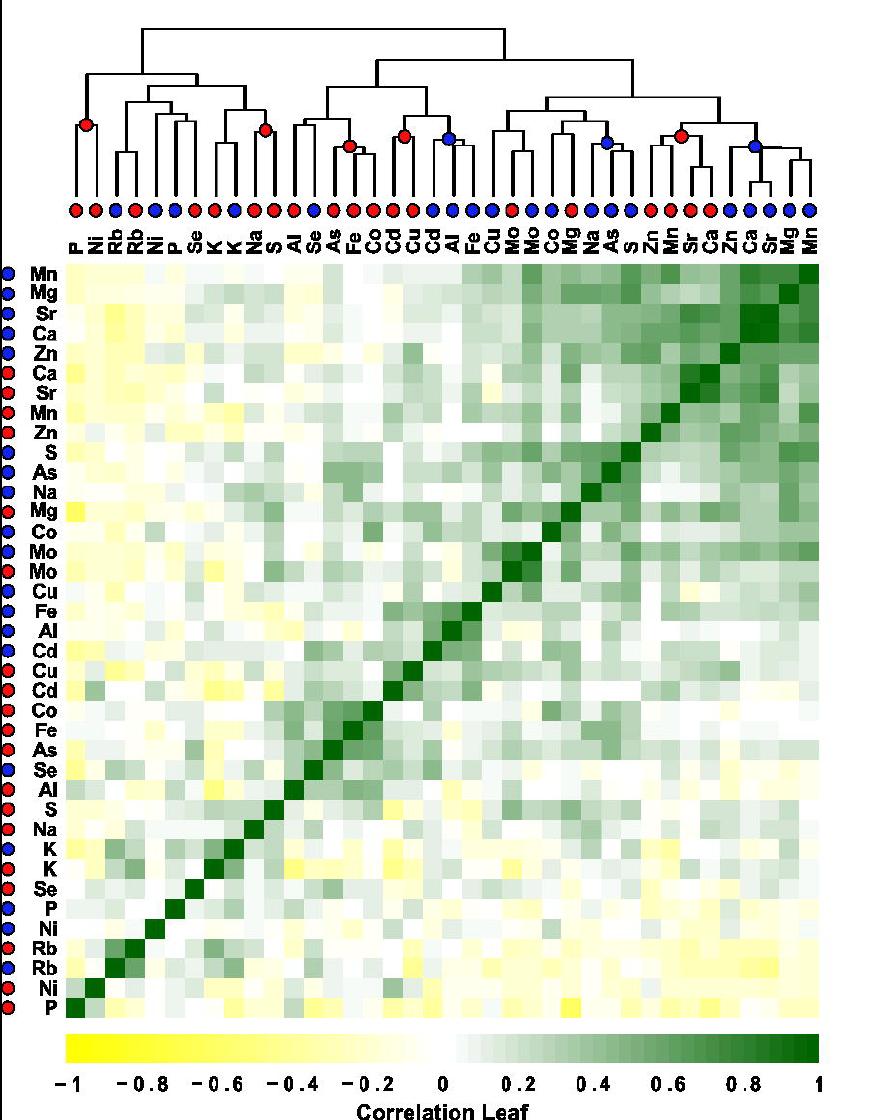
Patterns of ion concentration in leaves shift following inoculation with *Funneliformis mosseae.* Pairwise correlation of ion concentration in the leaves of thirty maize varieties grown with or without inoculation with *F. mosseae*. Data represented as Fig. 2.

To further investigate patterns of covariance in ion concentration, a principal component (PC) analysis was performed. Root and leaf data were analyzed separately (Fig. 4; Table 2). In both tissues, NC and M plants were partially separated on the basis of the first two PCs (representing 48% and 40% of the total variance in roots and leaves, respectively). In roots, the ions with greatest representation in PC1 were Al, As, Co, Fe, Ni and Rb (combined contribution of 71%; Fig. 5). Al, As, Co, Fe, Ni and Rb were clustered in the covariance matrix, and, as is consistent, contributed similarly to the PCs (Fig. 4; Table 2), their concentration generally decreasing in M plants. A significant reduction in Co and Ni was observed also in the single ion analysis, while median root concentrations of Al, As and Fe were reduced in M plants (Fig. 1; Table 1). Ca, Na, Mo, S, Sr and Rb contributed most to root PC2 (79% total; Fig.5), largely increasing in concentration in M plants, again consistent with the single ion analysis (Fig. 1; Table 1). Rb contributed equally to both PC1 and PC2 (7.8% and 5.9%, respectively) and showed no clear pattern with respect to inoculation (Fig. 1, Fig. 5). In leaves, PC1 and PC2 were predominantly represented by Ca, Mg, Mn, Mo, Sr and S (70%), and Al, As, Co, Fe, Ni and P (74%), respectively (Fig. 5; Table 2). NC and M treatments were best distinguished by PC2, generalized by an increase in the leaf concentration of Al, As, Co, Fe, Ni and P in M plants that was consistent with the single ion analysis (Fig. 1; Table 1). The contribution of P to the PCs was notably low, and in a direction opposite to the ions making the greatest contributions (Fig. 4), reflecting the negative correlations observed between concentrations of P and these ions, not only between treatments, but among genotypes within a single treatment (Fig. 3, S1, S2).

**Fig. 4.**
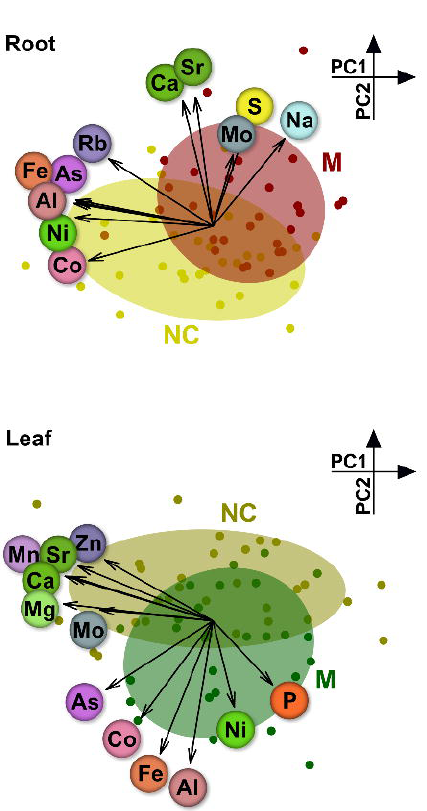
Inoculation with *Funneliformis mosseae* impacts the root and leaf ionomes. Differentiation of the mycorrhizal and non-mycorrhizal ionome. Principal component (PC) analysis of the concentration of nineteen ions in the roots and leaves of the thirty maize varieties (points) grown with (M) or without (NC) inoculation with *F. mosseae.* Biplot showing scores in the first two principal components (PC1: x-axis, PC2: y-axis). The sign and magnitude of the contribution of selected ions is shown by arrows. Ions are shown using conventional element coloring.

**Fig. 5.**
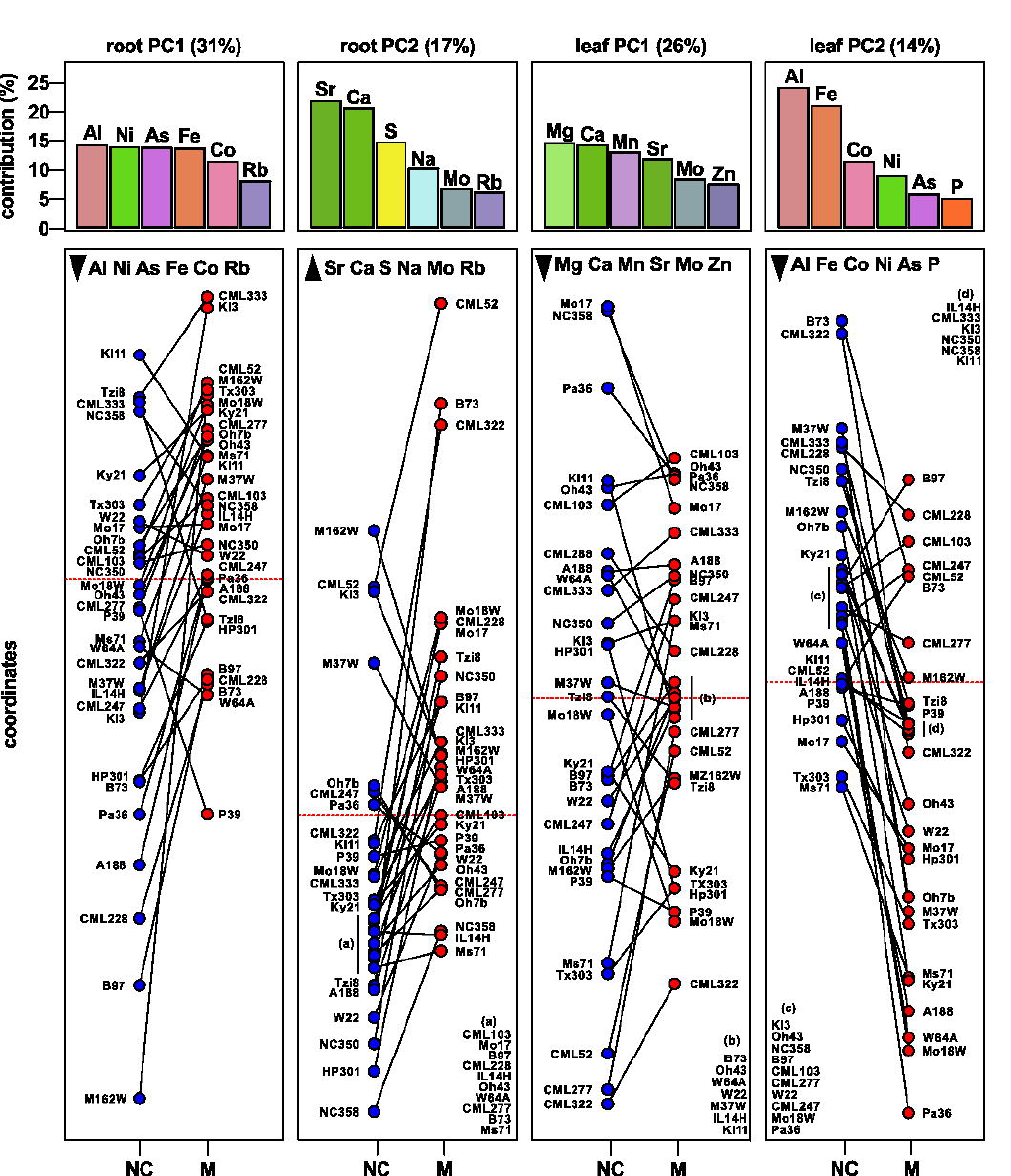
Ionome responses to inoculation with *Funneliformis mosseae* are dependent on plant genotype. Principal component (PC) coordinates for thirty maize inbred lines, for the first two PCs in analysis of the root and leaf ionomes. For each PC, the upper panel indicates the total contribution of the PC, along with the contributions of the six most important ions to that PC. Bars are filled using conventional coloring. For each genotype, the coordinates in a given PC are shown for non-inoculated (NC, blue points) and inoculated (M, red points) plants, linked by a line segment indicating the reaction norm (a plot of phenotype against environment, here contrasting NC and M). Coordinate units are arbitrary and scaled differently in the four panels, zero indicated by a red dashed line. Arrowheads in the top left of each coordinate panel indicate the direction of increasing concentration of the associated ions with reference to the y axis. Lower case letters indicate tightly clustered groups of genotypes that could not be clearly labeled and that are consequently presented in the corners of the relevant panels.

**Table 2.** Principal component analysis of the concentration of nineteen ions in roots and leaves. Scores of ions on the first three principal components (PCs) in root and leaf analysis. Coordinates were scaled x10 and rounded to two decimal places. Var, the percentage of variance associated with each PC.

|  | Root |  |  | Leaf |  |  |
| --- | --- | --- | --- | --- | --- | --- |
|  | PC1 | PC2 | PC3 | PC1 | PC2 | PC3 |
| Var | 31% | 17% | 12% | 26% | 14% | 10% |
| Al27 | -9.12 | 1.48 | -2.6 | -1.28 | -8.01 | -0.47 |
| As75 | -8.99 | 1.71 | -1.83 | -6 | -3.88 | 1.27 |
| Ca43 | -1.97 | 8.21 | -0.66 | -8.32 | 2.5 | -0.94 |
| Cd111 | -5.3 | -1.65 | 7.17 | -2.53 | -2.06 | -2.79 |
| Co59 | -8.16 | -2.26 | 3.5 | -4.05 | -5.48 | 0.78 |
| Cu65 | -4.16 | -3.04 | -0.53 | -3.74 | -3.16 | -2.43 |
| Fe57 | -8.93 | 1.56 | -3.19 | -2.97 | -7.47 | 0.38 |
| K39 | 2.61 | 2.74 | 2.54 | 0.81 | 3.44 | 7.28 |
| Mg25 | -1.87 | 3.24 | 2.86 | -8.41 | 0.95 | 2.04 |
| Mn55 | -6 | -1.47 | 5.84 | -7.96 | 2.48 | -0.67 |
| Mo98 | 1.33 | 4.67 | 5.95 | -6.35 | 0.63 | -0.71 |
| Na23 | 4.71 | 5.73 | -1.88 | -3.69 | -1.69 | 5.23 |
| Ni60 | -9.02 | 0.58 | 0.7 | 1.2 | -4.84 | -0.97 |
| P31 | -3.13 | 2.8 | -1.27 | 3.36 | -3.59 | 1.75 |
| Rb85 | -6.85 | 4.42 | -3.9 | 3.59 | -0.04 | 6.23 |
| S34 | 2.35 | 6.91 | 4.04 | -5.88 | -1.79 | 5.81 |
| Se82 | -1.37 | 1.73 | 3.88 | 0.47 | -0.15 | 1.38 |
| Sr88 | -1.14 | 8.45 | -0.41 | -7.58 | 3.08 | -1.57 |
| Zn66 | -4.42 | -3.64 | -0.01 | -6.03 | 3.4 | -0.43 |

### Differences in the ionome response to AM inoculation among maize varieties indicates the importance of host genotype

To investigate the importance of plant genotype to the response to inoculation with *F. mosseae,* the change in ionomic PC scores between NC and M treatments was compared (Fig. 5). Reaction norm plots reflected the main effect of inoculation in roots and shoots *(c.f.* Fig.1), but revealed also evidence of genotype specific differences (indicated by non-parallel lines in Fig. 5). As observed in PC biplots (Fig. 4), the difference between NC and M treatments was best captured by PC2, in both roots and leaves. In roots, there was a clear trend towards an increased PC2 score in inoculated plants, related to increasing concentrations of Ca, Na, Mo, S, Sr and Rb; in leaves the trend was towards a lower score in PC2, related to increasing concentrations of Al, As, Co, Fe, Ni and P. A number of lines, however, did not follow these general trends: in roots CML247, Ki3, M162W, M37W, Oh7b and Pa36 showed a reduction in PC2 when inoculated; in leaves, the lines B97, CML52, CML103 and CML247 showed an increase in PC2 when inoculated. These lines were not found to be exceptional with regard to growth response in the previous analysis (Sawers *et al.*, 2017), although it should be noted that P, the limiting nutrient, made only minor contributions to these PCs. As would be predicted by the clustering results (Fig. 2, 3), there were instances in which genotype specific effects were correlated among ions, well illustrated by the behavior of Al and Fe in the roots (Fig. 6).

**Fig. 6.**
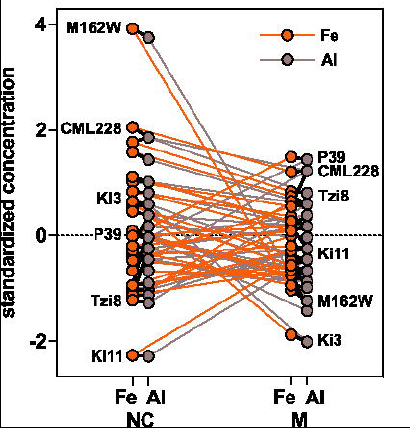
Root Fe and Al responses to AM colonization are correlated across genotypes. Standardized concentrations (z score) of Fe and Al in the roots of thirty maize varieties, grown with (M) or without (NC) inoculation with *Funneliformis mosseae.* The reaction norm (plot of phenotype against environment, here contrasting NC and M) for each ion-genotype combination is shown by a colored line. Concentrations of the two ions for each genotype within a treatment are connected by a black line. Selected varieties are labeled.

## DISCUSSION

Measurement of the concentration of nineteen ions in the leaves and roots of maize seedlings grown with or without inoculation with the AM fungus *Funneliformis mosseae* revealed coordinated changes in response to AM colonization. By using a panel of maize varieties designed to maximize genetic diversity, it was possible to both make meaningful generalizations and to examine patterns of covariation in ion concentration. Analysis of single ions and PCA indicated AM colonization to be associated with an increase in the root concentration of Ca, Na, Mo, P, Rb, S and Sr, and a decrease in Cd, Co, Cu, Mn, Ni and Zn (Fig. 1, 4, 5; Table 1). In the leaves, similar analyses revealed an increase in Al, As, Co, Fe, Na, Ni and P, and decrease of Mn and Zn in colonized plants (Fig. 1, 4, 5; Table 1). In single ion analysis, the concentrations of Mn, Na, Ni, P and Zn responded (p < 0.05) similarly to AM colonization in both roots and leaves, with Ni the only ion for which the sign of the response differed between the two tissues (Fig. 1; Table 1). Given that the analysis cannot distinguish between elements within the root and elements adhering to the root surface, the overlap between root and shoot provides an important indication that the changes observed in the root reflect differences in uptake. Although leaf P concentrations were at deficient levels (Reuter and Robinson, 1997), and variation in P concentration was shown previously to correlate well with plant growth in this experiment (Sawers *et al.*, 2017), the P response to AM inoculation was far from the most significant change to the ionome, nor did P contribute greatly to PCs (Fig. 1, 5; Table 1).

One of the most significant (p < 0.001) changes in the ionome of AM plants was the reduction in concentration in roots of Co and the potentially toxic, non-essential heavy metal Cd (Fig. 1; Table1). Although Co and Cd concentrations were relatively low (Reuter and Robinson, 1997) in both NC and M plants, these data are consistent with the previously reported role of AM fungi in protecting the host plant from accumulation of toxic elements (Göhre and Paszkowski, 2006). Mn concentration was also reduced, although non-limiting, in M plants, in both roots and leaves (Fig. 1; Table 1). Reduced Mn accumulation in M plants has been reported previously in maize and other plants, and attributed to reduced plant production of P-mobilizing carboxylates *(e.g.* Kothari *et al.*, 1991; Posta et al., 1994; Nazeri *et al.*, 2013; Gerlach *et al.*, 2015). Genotypes varied in Mn accumulation, and a negative correlation ( r = -0.35, p = 0.06; Fig. S1) was observed between P and Mn accumulation in the leaves of M plants. Measurement of the leaf Mn concentration has been proposed previously as a method to distinguish different strategies of P acquisition at higher taxonomic levels: concentrations of Mn will tend to be higher in species that exude carboxylate to favor direct P acquisition, due to increased Mn mobility in the rhizosphere when compared with species that tend to acquire P via mycorrhizae (Lambers *et al.*, 2015). Here, an intraspecific correlation was observed between Mn and P concentrations in M plants that might be exploited in the evaluation and genetic mapping of P uptake strategies. Correlations were observed also between P and Mg concentrations in M plants: a negative correlation in leaves (r = -0.62, p < 0.01; Fig. S2) and a positive correlation (r = 0.47, p < 0.01; Fig. S2) in roots. Cluster analysis of pairwise correlations revealed further groups of elements that varied together across genotypes (Fig. 2, 3). A number of clusters were common to NC and M plants, although the correlations between the two treatments were not strong, indicating a coordinated but genotype specific response: *i.e.* responses differed, but for a given genotype, the ions in a cluster responded in a similar way, a pattern well illustrated by concentrations of Al and Fe in the roots (Fig. 6)

In contrast to controlled experimental systems, field soils present a far more complex range of physicochemical properties, promoting correlation and interaction in the availability of nutrients to plants. For example, P deficiency is often accompanied by insufficiency of other nutrients, such as Ca, Mg and Zn in acid soils, or Fe and Zn in alkaline conditions (Calderón-Vázquez *et al.*, 2009; Hinsinger, 2001; Osaki *et al.*, 1999; Uexküll and Mutert, 1995). Soil water content will have nutrient specific effects: uptake of K and P is strongly reduced at low soil water content; uptake of Ca and Mg is less effected by water content (Talha *et al.*, 1979). It is clear that AM symbiosis has a broad impact on the host ionome beyond enhancement of P uptake. Although the direct fungal contribution to the uptake of each nutrient was not quantified, genotype specific responses are consistent with variation in symbiotic function, as reported previously with respect to P (Sawers *et al.*, 2017). Strong genotype specific responses were observed also for a number of elements which showed no change in mean concentration between NC and M plants when the panel was considered as a whole (e.g. Al and Fe in roots; Fig. 6), suggesting AM symbiosis to indeed impact, directly or indirectly, uptake of such elements. An analogy can be made to so-called “hidden” P uptake, the significant substitution of AM mediated for direct P uptake that can occur in colonized plants, even if the final P concentration, or indeed plant performance, are equivalent to non-colonized levels (Smith *et al.*, 2003).

Successful application of AM fungi in agricultural systems requires a suitable combination of plant host genotype, fungal community, soil environment and management (Fester and Sawers, 2012). It remains to be seen to what degree such a complex system can be manipulated, or, ultimately, optimized. AM symbiosis, however, represents far more than a simple exchange of carbon for P (Smith *et al.*, 2009). Indeed, AM symbiosis can be considered as a particular environmental modification that impacts all aspects of plant growth and development. Data presented here reveal both the profound impact of AM symbiosis on plant nutrition, and the importance of host genotype to the symbiotic outcome. Evaluation of the ionome is relatively stable and inexpensive, and might be readily incorporated into the evaluation of AM response in the field. Here, plants were grown in a greenhouse system under P limiting conditions, with other nutrients supplied in excess, and, as is consistent, no clear pattern was observed between the concentration of elements other than P and plant growth. Under a specific set of field conditions, however, the broad impact of AM symbiosis on the ionome beyond P acquisition may indeed impact performance, through to final yield and the nutritional quality of the grain.

## MATERIALS AND METHODS

### Growth of maize diversity panel inoculated with *Funneliformis mosseae*

As described previously (Sawers *et al.*, 2017), a panel of 30 diverse maize lines, comprising the 26 diverse inbred founders of the maize NAM population (McMullen *et al.*, 2009), Pa36 (a line tolerant of low P availability; Kaeppler *et al.*, 2000), and the broadly used reference lines B73 and W22, and W64A (a line used previously for study of AM symbiosis; Paszkowski *et al.*, 2006), was evaluated with (M) or without (NC) inoculation with *F. mosseae* (isolate number 12, European Bank of Glomales, http://www.kent.ac.uk/bio/beg/), as previously described (Sawers *et al.*, 2017). Briefly, plants were grown in 1 L pots, in sand/clay (9:1 vol/vol), fertilized three times per week with 100 ml of modified Hoagland solution (Hoagland and Broyer, 1936) containing 10% (100μM) of the standard concentration of KH_2_PO_4_, the potassium concentration being maintained by addition of KCl. A total of 1200 plants (30 genotypes x 2 treatments x 6 replicates) were grown in a complete block design, over five separate plantings, at the University of Lausanne, Switzerland. Plants were harvested after 8 weeks and shoot dry weight measured (SDW). For two plantings (corresponding to six complete blocks) roots were collected also, and stained to confirm efficacy of the fungal inoculum as previously described (Gutjahr *et al.*, 2008).

### Determination of elemental concentration by ICP-MS analysis

Root and shoot samples were analyzed by inductively coupled plasma mass spectrometry (ICP-MS) to determine the concentration of twenty metal ions. Weighed tissue samples were digested in 2.5mL concentrated nitric acid (AR Select Grade, VWR) with an added internal standard (20ppb In, BDH Aristar Plus). Sample digestion and dilution was carried out as described previously (Ziegler *et al,*2013). Concentration of the elements B11, Na23, Mg25, Al27, P31, S34, K39, Ca43, Mn55, Fe57, Co59, Ni60, Cu65, Zn66, As75, Se82, Rb85, Sr88, Mo98 and Cd111 was measured using an Elan 6000 DRC-e mass spectrometer (Perkin-Elmer SCIEX) connected to a PFA microflow nebulizer (Elemental Scientific) and Apex HF desolvator (Elemental Scientific). A control solution was run every tenth sample to correct for machine drift both during a single run and between runs. Measurements for B11 were not considered further as concentrations were apparently below the level of reliable detection. The ICP-MS technique does not allow determination of N concentration.

### Statistical analysis

Least squares (LS) means (lsmeans::lsmeans; Lenth, 2016) for concentrations were obtained based on fixed effect model for each genotype x ion x inoculation x tissue (root or shoot) combination. Differences in measured traits between treatments were investigated by Wilcoxon test (stats::wilcox.test; R Core Team, 2016) for paired comparisons, and by ANOVA (stats::lm) and *post hoc* Tukey HSD test (agricolae::test.HSD, de Mendiburu, 2016) for multiple comparisons. Percentage root length colonization data was square root transformed prior to analysis. Ion concentration LS means were used to calculate separate correlation matrices for root and shoot (Hmisc::rcorr; Harrel, 2016) that were visualized as heatmaps (gplots::heatmap.2, Warnes *et al.*, 2016), using the default hierarchical clustering. Principal component analysis (PCA) was performed separately for root and leaf samples, using all ion concentrations, with R statistics ade4::dudi.pca (Dray & Dufour, 2007) using centered and scaled data, and the results visualized with ade4::scatter. Only selected ions were included in the biplot.

## ACKNOWLEDGEMENTS

We thank Matthias Mueller for assistance with plant growth and evaluation.

## Supplementary Figures

**Fig. S1. P and Mn concentrations in plants with and without inoculation with *Funneliformis mosseae***. A - D concentrations (ppm) of P31 and Mn55 in the roots and leaves of non-colonized (NC) plants and plants inoculated with F. mosseae. Points represent mean values for each of thirty maize varieties.

**Fig. S2. P and Mg concentrations in plants with and without inoculation with *Funneliformis mosseae***. A - D concentrations (ppm) of P31 and Mg25 in the roots and leaves of non-colonized (NC) plants and plants inoculated with F. mosseae. Points represent mean values for each of thirty maize varieties.

## REFERENCES

Aibara, I. and Miwa, K. (2014) Strategies for optimization of mineral nutrient transport in plants: multilevel regulation of nutrient-dependent dynamics of root architecture and transporter activity. Plant Cell Physiol. 55: 2027–2036.

Bago, B., Pfeffer, P.E., Abubaker, J., Jun, J., Allen, J.W., Brouillette, J., et al. (2003) Carbon export from arbuscular mycorrhizal roots involves the translocation of carbohydrate as well as lipid. Plant Physiol. 131: 1496–1507.

Bates, D., Maechler, M., Bolker, B. and Walker, S. (2015) Fitting linear mixed-effects models using lme4. J. Stat. Software 67: 1–48. Accessed January 2017.

Baxter, I.R., Vitek, O., Lahner, B., Muthukumar, B., Borghi, M., Morrissey, J., et al. (2008) The leaf ionome as a multivariable system to detect a plant’s physiological status. Proc Natl Acad Sci USA 105: 12081–6.

Baxter, I.R. (2015) Should we treat the ionome as a combination of individual elements, or should we be deriving novel combined traits? J Exp Bot 66: 2127–31.

Bucher, M. (2007) Functional biology of plant phosphate uptake at root and mycorrhiza interfaces. New Phytol. 173: 11–26.

Calderón-Vázquez, C., Alatorre-Cobos, F., Simpson-Williamson, J. and Herrera-Estrella, L. (2009) Maize under phosphate limitation. In Handbook of Maize: Its Biology. Edited by Bennetzen, J. L. and Hake, S.C. pp. 381–404. Springer, New York.

Dray, S. and Dufour, A.B. (2007) The ade4 package: implementing the duality diagram for ecologists. J Statistical Softw. 22: 1–20.

Fester, T. and Sawers, R.J.H. (2011) Progress and challenges in agricultural applications of arbuscular mycorrhizal fungi. Crit Rev Plant Sci 30: 459–470.

Finlay, R.D. (2008) Ecological aspects of mycorrhizal symbiosis: with special emphasis on the functional diversity of interactions involving the extraradical mycelium. J. Exp. Bot. 59(5), 1115–1126.

Gerlach, N., Schmitz, J., Polatajko, A., Schlüter, U., Fahnenstich, H., Witt, S., et al. (2015) An integrated functional approach to dissect systemic responses in maize to arbuscular mycorrhizal symbiosis. Plant, cell & environment 38: 1591–612.

Giovannetti, M., Tolosano, M., Volpe, V., Kopriva, S. and Bonfante P. (2014) Identification and functional characterization of a sulfate transporter induced by both sulfur starvation and mycorrhiza formation in *Lotus japonicus*. New Phyt. 204: 609–619.

Göhre, V. and Paszkowski, U. (2006) Contribution of the arbuscular mycorrhizal symbiosis to heavy metal phytoremediation. Planta 223: 1115–1122.

Govindarajulu, M., Pfeffer, P.E., Jin, H., Abubaker, J., Douds, D.D., Allen, J.W., et al. (2005) Nitrogen transfer in the arbuscular mycorrhizal symbiosis. Nature 435: 819–823.

Guether, M., Neuhäuser, B., Balestrini, R., Dynowski, M., Ludewig, U. and Bonfante, P. (2009) A mycorrhizal-specific ammonium transporter from *Lotus japonicus* acquires nitrogen released by arbuscular mycorrhizal fungi. Plant Physiol. 150: 73–83.

Gutjahr, C., Banba, M., Croset, V., An, K., Miyao, A., An, G., et al. (2008) Arbuscular mycorrhiza-specific signaling in rice transcends the common symbiosis signaling pathway. Plant Cell 20: 2989–3005.

Harrell, F.E. (2016) Hmisc: Harrell Miscellaneous. R package version 3.17–4. https://CRAN.R-project.org/package=Hmisc. Accessed January 2017.

Harrison, M.J., Dewbre, G.R. and Liu, J. (2002) A phosphate transporter from *Medicago truncatula* involved in the acquisition of phosphate released by arbuscular mycorrhizal fungi. Plant Cell 14: 2413–2429.

Hetrick, Hetrick.B.A.,Wilson, Wilson.G.W. and Cox, T.S. (1992) Mycorrhizal dependence of modern wheat varieties, landraces, and ancestors. Can. J. Bot. 70: 2032–2040.

Hinsinger, P. (2001) Bioavailability of soil inorganic P in the rhizosphere as affected by root-induced chemical changes: a review. Plant and Soil 237: 173–195.

Hoagland, D.R. and Broyer. T.C. (1936) General nature of the process of salt accumulation by roots with description of experimental methods. Plant Physiol. 11: 451–507.

Jin, H., Pfeffer, P.E., Douds, D.D., Piotrowski, E., Lammers, P.J. and Shachar-Hill, Y. (2005) The uptake, metabolism, transport and transfer of nitrogen in an arbuscular mycorrhizal symbiosis. New Phytol. 168: 687–696.

Kaeppler, S.M., Parke, J.L., Mueller, S.M., Senior, L., Stuber, C. and Tracy, W.F. (2000) Variation among maize inbred lines and detection of quantitative trait loci for growth at low phosphorous and responsiveness to arbuscular mycorrhizal fungi. Crop Sci 40: 358–364.

Koegel, S., Lahmidi, N.A., Arnould, C., Chatagnier, O., Walder, F., Ineichen, K., et al. (2013) The family of ammonium transporters (AMT) in *Sorghum bicolor*; two AMT members are induced locally, but not systemically in roots colonized by arbuscular mycorrhizal fungi. New Phytol. 198: 853–865.

Koide, R.T., Li, M., Lewis, J. and Irby, C. (1988) Role of mycorrhizal infection in the growth and reproduction of wild vs. cultivated plants. Oecologia 77: 537–543.

Kothari, S.K., Marschner, H., and Romheld, V. (1991) Effect of a vesicular-arbuscular mycorrhizal fungus and rhizosphere micro-organisms on manganese reduction in the rhizosphere and manganese concentrations in maize (*Zea mays L.*). New Phytol. 117: 649–655.

Lambers, H., Hayes, P.E., Laliberté, E., Oliveira, R.S. and Turner, B.L. (2015) Leaf manganese accumulation and phosphorus-acquisition efficiency. Trends in Plant Science 20: 83–90.

Lenth, R. (2016) Least-squares means: the R package lsmeans. J. Stat. Software 69: 1–33. Accessed January 2017.

Liu, F., Xu, Y., Jiang, H., Jiang, C., Du, Y., Gong, C., et al. (2016) Systematic identification, evolution and expression analysis of the *Zea mays PHT1* gene family reveals several new members involved in root colonization by arbuscular mycorrhizal fungi. Int J Mol Sci 17: 10.3390/ijms17060930

Lynch, J. (1995) Root Architecture and Plant Productivity. Plant Physiol. 109: 7–13.

Marschner, P. (Ed.) (2012) Mineral Nutrition of Higher Plants. Academic Press, San Diego.

McMullen, M.D., Kresovich, S., Villeda, H.S., Bradbury, P., Li, H., Sun, Q., et al. (2009) Genetic properties of the maize nested association mapping population. Science 325: 737–40.

de Mendiburu, F. (2016) Agricolae: Statistical Procedures for Agricultural Research. R package version 1. 2–4. https://CRAN.R-project.org/package=agricolae. Accessed January 2017.

Nazeri, N. K., Lambers, H., Tibbett, M. and Ryan, M.H. (2013) Do arbuscular mycorrhizas or heterotrophic soil microbes contribute toward plant acquisition of a pulse of mineral phosphate? Plant and Soil. 373: 699–710.

Nadal, M.,Sawers, R.J.H., Naseem, S., Bassin, B., Kulicke, C., Sharman, A., et al. (2017) An N-acetylglucosamine transporter required for arbuscular mycorrhizal symbioses in rice and maize. Nat Plants 26: 17073.

Osaki, M., Friesen, D.J. and Rao, I.M. (1999) Plant adaptation to phosphorus-limited tropical soils. In Handbook of Plant and Crop Stress (Second Edition) pp. 61–95. CRC Press.

Parniske, M. (2008) Arbuscular mycorrhiza: the mother of plant root endosymbioses. Nat. Rev. Microbiol. 6: 763–775.

Paszkowski, U., Kroken, S., Roux, C. and Briggs, S.P. (2002) Rice phosphate transporters include an evolutionarily divergent gene specifically activated in arbuscular mycorrhizal symbiosis. Proc. Natl. Acad. Sci. USA 99: 13324–13329.

Paszkowski, U., Jakovleva, L. and Boller T. (2006) Maize mutants affected at distinct stages of the arbuscular mycorrhizal symbiosis. Plant J. 47: 165–173.

Porcel, R., Aroca, R., Azcon, R. and Ruiz-Lozano, J. M. (2016) Regulation of cation transporter genes by the arbuscular mycorrhizal symbiosis in rice plants subjected to salinity suggests improved salt tolerance due to reduced Na^+^ root-to-shoot distribution. Mycorrhiza 26: 673–684.

Posta, K., Marschner, H., Römheld, V. (1994) Manganese reduction in the rhizosphere of mycorrhizal and nonmycorrhizal maize. Mycorrhiza 5: 119–124.

R Core Team. (2016) R: A language and environment for statistical computing. R Foundation for Statistical Computing, Vienna, Austria. Version 3.3.1. URL https://www.R-project.org/. Accessed January 2016.

Reuter, D. and Robinson, J.B. (1997) Plant Analysis: An Interpretation Manual. Csiro Publishing.

Sawer, R.J.H., Gutjahr, C. and Paszkowski, U. (2008) Cereal mycorrhiza: an ancient symbiosis in modern agriculture. Trends Plant Sci 13: 93–97.

Sawers, R.J.H., Svane, S.F., Quan, C., Grønlund, M., Wozniak, B., González-Muñoz, E., et al. (2017) Phosphorus acquisition efficiency in arbuscular mycorrhizal maize is correlated with the abundance of root-external hyphae and the accumulation of transcripts encoding PHT1 phosphate transporters. New Phytol. 214: 632–643.

Schüpler, A., Schwarzott, D., and Walker, C. (2001). A new fungal phylum, the Glomeromycota: phylogeny and evolution. Mycological Research 105: 413–1421.

Smith, S.E. and Read D.J. (2008) Mycorrhizal symbiosis. Cambridge, UK: Academic Press.

Smith, S.E., Smith, F.A, and Jakobsen, I. (2003) Mycorrhizal fungi can dominate phosphate supply to plants irrespective of growth responses. Plant Physiol. 133: 16–20.

Smith, F.A., Grace, E.J. and Smith, S.E. (2009) More than a carbon economy: nutrient trade and ecological sustainability in facultative arbuscular mycorrhizal symbioses. New Phytol. 182: 347–58.

von Uexküll, H.R. and Mutert, E. (1995) Global extent, development and economic impact of acid soils. Plant and Soil 171: 1–15.

Vance, C.P. (2014) Symbiotic nitrogen fixation and phosphorus acquisition. Plant nutrition in a world of declining renewable resources. Plant. Physiol. 127: 390–397.

Warnes, G.R., Bolker, B., Bonebakker, L., Gentleman, R., Huber, W., Liaw, S., et al. (2016) gplots: Various R programming tools for plotting data. R package version 3.0.1. https://CRAN.R-project.org/package=gplots. Accessed January 2016.

Whitcomb, S. J., Heyneke, E., Aarabi, F., Watanabe, M. and Hoefgen, R. (2014) Mineral nutrient depletion affects plant development and crop yield. In Nutrient use efficiency in plants Edited by M. J. Hawkesford, M.J., Kopriva, S. and Kok, L. J. D. pp. 205–208. Springer International Publishing.

Willmann, M., Gerlach, N., Buer, B., Polatajko, A., Nagy, R., Koebke, E., et al. (2013) Mycorrhizal phosphate uptake pathway in maize: vital for growth and cob development on nutrient poor agricultural and greenhouse soils. Front Plant Sci 4: 533.

Yang, S.Y., Grønlund, M., Jakobsen, I., Grotemeyer, M.S., Rentsch, D., Miyao, A., et al. (2012) Nonredundant regulation of rice arbuscular mycorrhizal symbiosis by two members of the *PHOSPHATE TRANSPORTER1* gene family. Plant Cell 24: 4236–4251.

Ziegler, G., Terauchi, A., Becker, A., Armstrong, P., Hudson, K. and Baxter, I. (2013) Ionomic screening of field-grown soybean identifies mutants with altered seed elemental composition. Plant Genome 6: 1–9.

